# Climate change has already reshaped North American forest pest dynamics: Insights from multidecadal process-based modelling

**DOI:** 10.1101/2025.10.15.682655

**Authors:** Yan Boulanger, Chris J. K. MacQuarrie, Véronique Martel, Jacques Régnière, Rémi Saint-Amant, Kishan Sambaraju

**Affiliations:** Natural Resources Canada, Canadian Forest Service, Laurentian Forestry Centre, 1055 rue du Peps, Québec City, QC, G1V 4C7, Canada; Natural Resources Canada, Canadian Forest Service, Great Lakes Forestry Centre, 1219 Queen Street East Sault Ste. Marie, ON P6A 2E5 Canada

**Keywords:** ERA5, mountain pine beetle, spruce budworm, hemlock woolly adelgid, spongy moth, emerald ash borer, spruce beetle, southern pine beetle, temperature

## Abstract

Ongoing anthropogenic climate change has the potential to modify the population dynamics of forest pest insects by shifting the distribution of suitable climate conditions for their development. We used processed-based temperature-driven physiological models to assess the impact of changing climate conditions between 1951 and 2022 across North America on eight important forest pest insects, namely western spruce budworm, eastern spruce budworm, spongy moth, hemlock woolly adelgid, mountain pine beetle, southern pine beetle, spruce beetle and emerald ash borer. Our analyses revealed substantial changes in climate suitability resulting in pronounced northward and elevational shifts for most pest species although the magnitude and spatial patterns of these shifts varied both geographically and among species. We also showed that shifts in highly suitable conditions were more important at the colder edge (either northern or upper elevation) than at the warmer boundaries for several species, either driven by the Arctic amplification or elevation-dependant warming. Our results indicated that both the total area and the host tree biomass exposed to highly suitable climate conditions have increased for many pest species over the last decades, further exposing ecosystems to elevated risk. Our analyses also identified areas (e.g., western Canada) that are increasingly exposed to overlapping, potentially cumulative and interacting, biological disturbances. We also showed that climate change has already contributed to increasing the climatic suitability and geographic spread of exotic forest pest species in North America. Changes in climate suitability over the past seven decades across North America likely represent early signals of continued and potentially accelerating shifts under ongoing anthropogenic climate forcing. In this context, efforts to limit the expansion of pest populations under future climate change scenarios will be key to mitigating their cultural, ecological and economic impacts.

## Introduction

Forest insect pests are the most significant disturbance affecting North American forests (Dukes et al. 2009; Cooke et al. 2021). These disturbances occur across a wide range of forest biomes, including temperate, and boreal forests, highlighting their broad ecological relevance. During outbreak episodes, these species can cause extensive tree mortality, profoundly altering forest structure and composition. Such changes have cascading effects on ecosystem processes, notably affecting carbon and nutrient cycling (Dymond et al. 2010). Beyond ecological impacts, several species also cause substantial economic losses for the forest industry (Hennigar et al. 2011) and can severely degrade the ecological and aesthetic value of urban forests (Sydnor et al. 2007).

Climate plays a critical role in regulating the population dynamics of forest insect pests. As ectothermic organisms, the development, phenology, and survival of insects is highly sensitive to temperature and other climatic variables (Culos and Tyson 2014). Weather conditions influence egg hatching, larval growth rates, dispersal and mating opportunities, and overwintering survival (Williams and Liebhold 2002; Régnière et al. 2012). For example, warmer temperatures can accelerate insect development, increase fecundity, improve synchrony with host phenology, and enhance winter survival. In addition to these direct effects, climate also indirectly shapes insect population dynamics by altering the abundance and effectiveness of natural enemies (e.g., predators and parasitoids) (Seehausen et al. 2017, 2018; Régnière et al. 2021a) and by modifying host tree vulnerability (Kolb et al. 2016; Münro et al. 2021). As a result, ongoing anthropogenic climate change has been found to influence pest dynamics throughout the world’s forests. For instance, increasingly milder winter conditions have been associated with better winter survival in various pest species (Bale and Hayward 2010; Schneider et al. 2021) and earlier seasonal activity (Huang and Li 2015). These climate-driven changes have already contributed to increased severity and frequency in outbreak patterns (Jepsen et al. 2008; Pureswaran et al. 2015; Sambaraju et al. 2019). Additionally, greater synchrony between the timing of insect activity and host plant phenology with warming temperatures was shown to increase host susceptibility (Pureswaran et al. 2018; Ekholm et al. 2019), while disrupting predator-prey and parasitoid-host interactions (Stireman et al. 2005). Of particular concern is the growing body of evidence reporting poleward and altitudinal shifts or expansions in the geographic distribution of numerous insect pest species (Soja et al. 2007; Jepsen et al. 2008; Sambaraju et al. 2019; Ma et al. 2021). Such range expansions may subject previously unaffected tree species and forest biomass to increased pest pressure in regions where climatic conditions were historically unsuitable. This shift can undermine the resilience of forest ecosystems and challenge existing forest management practices that are not adapted to these emerging threats. As climate conditions continue to evolve, the frequency and intensity of damages from forest insect pests are expected to increase (Williams and Liebhold 1995; Fleming and Volney 1995; Logan et al. 2003), posing growing challenges for North American forest health and management (Marini et al. 2017).

The impacts of climate change on pest dynamics are inherently complex, as responses vary not only among species but also across seasons and bioclimatic regions (Robinet and Roques 2010). Although numerous studies have projected future impacts of climate change on pest distributions and population dynamics (e.g., Régnière et al. 2012), surprisingly few have examined multidecadal historical trends in climatic suitability for these species. Yet, such retrospective analyses are crucial for understanding how ongoing climate change is already influencing pest performance as well as for validating projections of future shifts. Moreover, while a few retrospective studies have provided valuable insights (e.g., Boulanger et al. 2025), most have focused on individual pest species in isolation (Régnière et al. 2019). A comprehensive, comparative analysis across multiple taxa and forest ecosystems remains largely absent, limiting our understanding of broader, cross-species, cumulative patterns and vulnerabilities under climate change. Furthermore, many large-scale studies on the impacts of climate change on forest pests have relied on correlative models such as species distribution models (e.g., De Santis et al. 2014; Liang and Fei 2014; Hill et al. 2016; Sidder et al. 2016; Cornelson et al. 2024). While useful for identifying potential shifts in species ranges, these models are limited in their ability to capture the complex, nonlinear responses of individual insect life stages to recent changes in climate. In contrast, mechanistic models that simulate the development of individuals and that are driven by weather inputs allow for a more detailed examination of how climate affects insect phenology, survival, and overall population dynamics (Lehmann et al. 2018). By explicitly representing biological processes, process-based ecophysiological models provide a more nuanced understanding of how forest pest dynamics may have shifted under recent climate change, offering insights that go beyond those obtainable through correlative approaches alone (Morin and Thuiller 2009).

In this study, we evaluate recent trends in climate suitability for several key forest insect pest species across North America. Specifically, we use published process-based ecophysiological models to estimate where, and to what extent, climate conditions conducive to pest development have changed between 1951 and 2022 for eight major species affecting important forest ecosystems in North America. We further investigate the extent to which suitable climate conditions have shifted northward or to higher elevations over time and assess how these shifts have increased the exposure of host tree biomass to pest risk.

## Methods

### Study overview

This study focuses on eight major North American forest insect species: spongy moth (SM, *Lymantria dispar* (L.) [Lepidoptera: Erebidae]; formerly ‘Gypsy moth’), western spruce budworm (WSBW, *Choristoneura occidentalis* Freeman [Lepidoptera: Tortricidae], = C*horistoneura freemanii* Razowski), eastern spruce budworm (ESBW, *Choristoneura fumiferana* (Clemens) [Lepidoptera: Tortricidae]), hemlock woolly adelgid (HWA, *Adelges tsugae* Annand [Hemiptera: Adelgidae]), mountain pine beetle (MPB, *Dendroctonus ponderosae* (Hopkins) [Coleoptera: Curculionidae]), southern pine beetle (SPB, *Dendroctonus frontalis* Zimmermann [Coleoptera: Curculionidae]), spruce beetle (SB, *Dendroctonus rufipennis* Kirby [Coleoptera: Curculionidae]), and emerald ash borer (EAB, *Agrilus planipennis* Fairmaire [Coleoptera: Buprestidae]). These species attack a wide array of host tree species and are distributed across a broad range of North American biomes, from the pine barrens of the southeastern United States to the boreal forests of northern Canada and Alaska. Among them, SM, EAB and HWA are non-native species, while the other five species are native to North America. All these species are known to cause extensive damage to host trees notably during outbreaks which can strongly impact forest succession or forest sector economics, or both.

We used published process-based, weather-driven ecophysiological models to evaluate weather-driven changes in annual species performance measures from 1950 to 2022 (Table 1). These models are essentially temperature-driven and simulate key stages of insect development, reproduction, and/or survival. These performance metrics vary by model and species because each species responds differently to temperature across critical life stages.

**Table 1.**
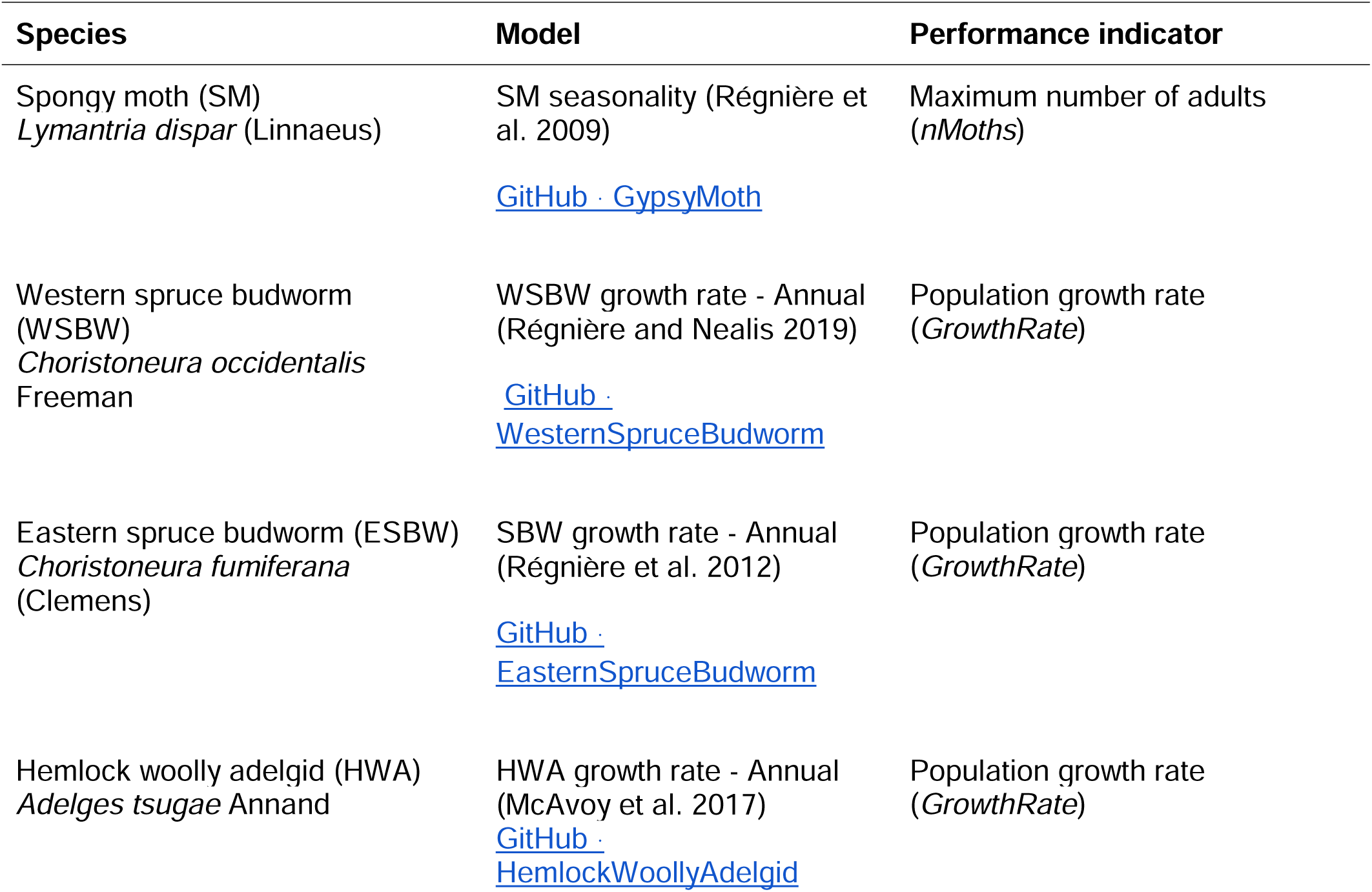
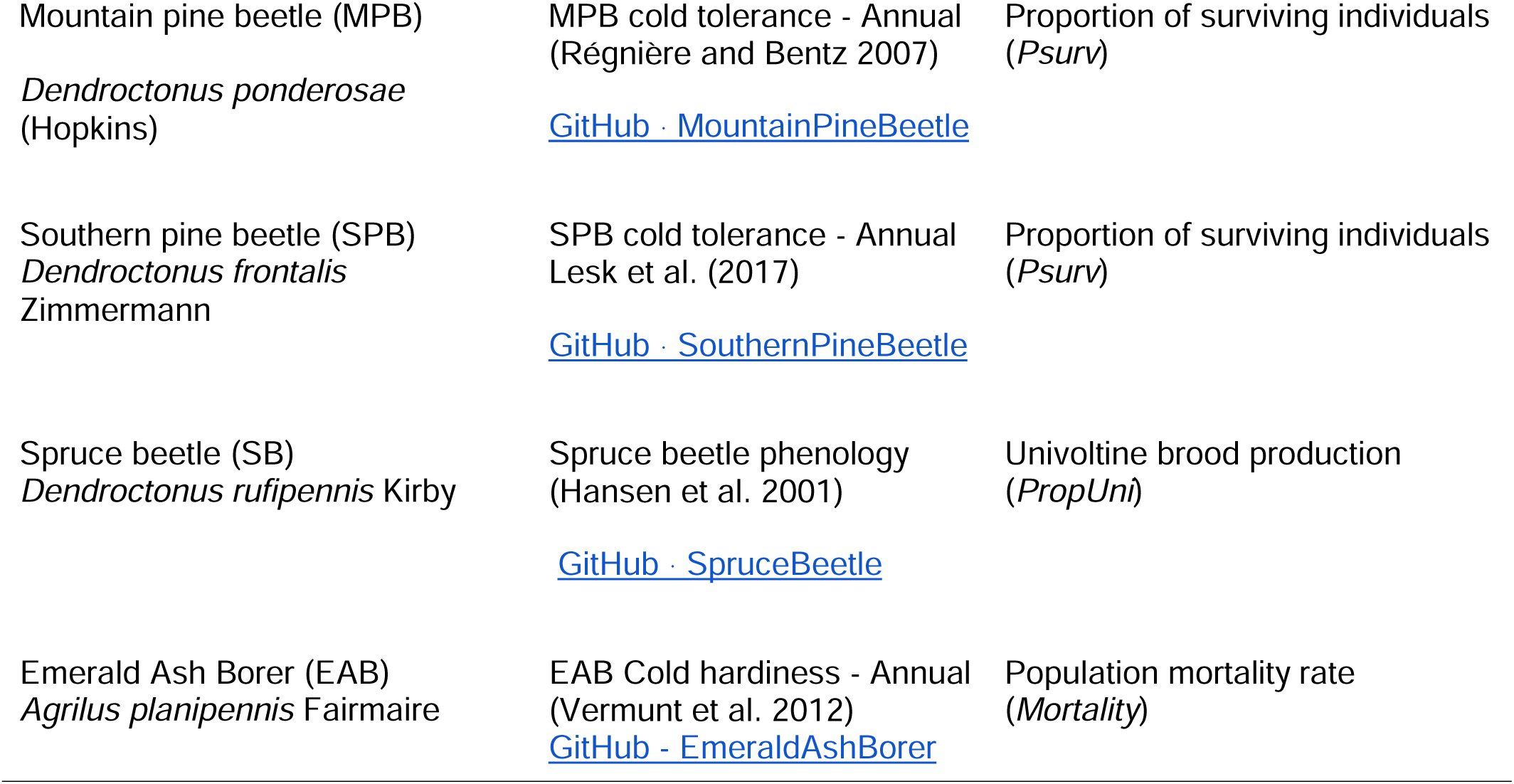
Species included in this study, along with the process-based ecophysiological models used for each species and the corresponding variables selected as performance indicators. All models are available from the GitHub link provided for each species.

Spatiotemporal trends in performance metrics and host tree biomass at risk were then assessed. Detailed information on each pest species and the corresponding ecophysiological model used to assess climate-driven performance is provided below.

### Spongy moth

Spongy moth (SM) is a non-native lepidopteran defoliator originating from Eurasia, first introduced to North America in 1869 near Boston, Massachusetts (Riley & Vasey 1870). Since its introduction, it has expanded its range across much of northeastern North America, including New England and eastern Canada. This species is characterized by periodic, synchronous outbreaks occurring every 10–15 years, typically lasting 1–3 years. It is highly polyphagous, feeding on over 500 plant species (Liebhold et al. 1995), although oaks (*Quercus* spp.) are particularly susceptible during outbreak events.

The model used to assess the performance of SM is based on the work of Régnière et al. (2009), who developed a temperature-driven phenology model to estimate the probability of individuals completing their life cycle under given climatic conditions. The model relies on daily temperature inputs and integrates developmental thresholds and timing based on earlier phenological models (Gray et al. 1991, 1995, 2001; Logan et al. 1991; Sheehan 1992; Régnière and Sharov 1998; Régnière and Nealis 2002). It simulates progression through the full life cycle, returning the proportion of the simulated population that successfully completes development within a single year under annual climate scenarios. From these outputs, we selected the maximum number of adult moths (*nMoths*) produced as the performance indicator.

### Western spruce budworm

Western spruce budworm (WSBW) is a native lepidopteran defoliator widely distributed across western North America, from British Columbia to the southwestern United States. It primarily feeds on Douglas-fir (*Pseudotsuga menziesii*), true firs (*Abies* spp.), and spruces (*Picea* spp.).

Outbreaks historically recurred every 30 to 40 years but since 1970, three major outbreaks have been recorded in western Canada, with evidence of expansion into more northerly and higher elevation areas (Régnière and Nealis 2019). Outbreaks can persist for several years, causing significant defoliation, growth loss, and tree mortality, especially in dense or multi-aged stands. This species is considered one of the most damaging defoliators in western North America’s coniferous forests (Nealis and Régnière 2021).

We used the population growth model developed by Régnière and Nealis (2019) to assess the performance of WSBW. This phenological, individual-based model simulates the ability of the species to complete its life cycle by tracking the number and physiological state of individuals at each developmental stage, from overwintering L2 larvae (L2o) through successive instars (L2– L6), pupae, adults, eggs, and the subsequent generation of L2o. The model is primarily temperature-driven and incorporates four key mortality factors: (1) phenological mismatch with host availability, (2) fecundity of surviving individuals, (3) frost exposure during the maturation and reproductive periods, and (4) energy depletion during overwintering. From these simulations, we calculated the simulated population growth rate (*GrowthRate*) which was used as the performance indicator for this pest.

### Eastern spruce budworm

Eastern spruce budworm (ESBW) is a native lepidopteran defoliator and one of the most destructive forest pests in eastern Canada and the northeastern United States (Williams and Liebhold 2000). It primarily feeds on balsam fir (*Abies balsamea*) and white spruce (*Picea glauca*), though other spruce species may be attacked. Outbreaks typically occur every 30–40 years and can last a decade or more, resulting in extensive tree mortality over large areas and having major ecological and economic consequences (Hennigar et al. 2011). Cold temperatures in the northern part of the species’ range limit its ability to complete development, while in the southern range, warmer conditions constrain populations by accelerating energy depletion during diapause (Régnière et al. 2012).

In our analyses ESBW climate-driven performance was assessed using an individual-based model (Régnière et al. 2012) that simulates development, survival, and reproduction from overwintering larvae. The model uses daily temperatures to simulate the development of all life stages (egg, L1, L2o, L2 to L6, pupa, and adult). It produces estimates of phenology, reproduction, overwinter survival, and potential annual population growth (*GrowthRate)*. This latter variable was used as a performance indicator. A complete description of the model is available in Régnière et al. (2012).

### Hemlock woolly adelgid

Hemlock woolly adelgid (HWA) is an invasive, sap-feeding insect native to East Asia (Limbu et al. 2018). It was first detected in the eastern United States in the 1950s and has since spread throughout a great portion of the range of eastern hemlock (*Tsuga canadensis*) and Carolina hemlock (*Tsuga caroliniana*) in the United States. It was detected in Ontario and Nova Scotia in the 2010s where host mortality has been reported (MacQuarrie et al. 2025b) Infestations cause tree decline and mortality over several years and its spread has had profound effects on forest structure and biodiversity in hemlock-dominated ecosystems (Trotter et al. 2013). Endemic, native populations of HWA are also found in the US Pacific northwest and British Columbia but their impacts in this region are negligible (Havill et al. 2016) and as such, only the eastern portion of its range was considered in our analyses.

Populations of HWA are particularly sensitive to cold temperatures during winter. In this study, we predicted population growth rates as a function of survival during winter based on models developed by McAvoy et al. (2017). These models predict winter mortality using three temperature-based metrics: the winter’s lowest minimum temperature (Tmin), the number of days with mean temperature below -1°C, and the mean temperature over the three days preceding Tmin to account for potential cold acclimation. We used the models’ estimated population density changes (population growth rate*, GrowthRate*) assessed from the overwintering (sistens) generation to the following summer (progrediens) generation as the performance indicator for HWA.

### Mountain pine beetle (*Dendroctonus ponderosae*)

Mountain pine beetle (MPB) is a bark beetle native to western North America and one of the most economically damaging forest insects on the continent (Dahr et al. 2016). It primarily attacks lodgepole pine (*Pinus contorta*) but also infests other pine species such as ponderosa pine (*P. ponderosa*) and jack pine (*P. banksiana*). Outbreaks are driven by climate conditions that favor beetle survival and development, particularly warmer winters and longer growing seasons (Sambaraju et al. 2019). A severe outbreak that began in the early 2000s caused extensive pine mortality across millions of hectares in British Columbia and resulted in an unprecedented range expansion of MPB east of the northern Rocky Mountains into the boreal forest of western Canada (Cooke and Carroll 2017).

Several process-based and hybrid models (Preisler et al. 2012, Sambaraju and Goodsman 2021) grounded in ecophysiological principles have been developed to characterize aspects of MPB biology, including winter survival, climate suitability, and adaptive seasonality (see Logan and Bentz 1999; Hicke et al. 2006; Bentz et al. 2010). Yet, winter survival is considered a critical factor influencing MPB population dynamics. Historically, extreme minimum temperatures in colder regions (i.e., higher elevations and northern latitudes) in western North America suppressed outbreaks, but recent warming has relaxed these limits (Buotte et al. 2016). Cold tolerance was also deemed one of the best determinants of outbreak potential (Preisler et al. 2012), with outbreak dynamics strongly influenced by the timing, frequency, and duration of cold snaps (Sambaraju et al. 2012). For this study, we selected the winter survival model developed by Régnière and Bentz (2007) to estimate annual survival probability (*Psurv*) as the performance indicator for MPB. The model uses a mechanistic framework developed to simulate the distribution of supercooling points in MPB by linking daily temperature variations to physiological processes acquisition and loss of cold tolerance. The model tracks individuals transitioning through three stages: (i) an active, feeding stage with no cold hardiness, (ii) a preparatory stage in which feeding ceases, the gut contents and ice-nucleating agents are eliminated, and (iii) a fully cold-hardy stage characterized by maximal accumulation of cryoprotectants (e.g., glycerol). Transitions through these stages is driven by opposing rates of cold acclimation and deacclimation in response to environmental cues.

### Southern pine beetle

Southern pine beetle (SPB) is a highly aggressive tree-killing bark beetle native to the southeastern United States (Pye et al. 2011). Under favorable conditions, populations can erupt into outbreaks that cause rapid and widespread tree mortality. It attacks a range of pine species, especially loblolly (*Pinus taeda*), shortleaf (*P. echinata*), and slash pine (*P. elliottii*) in the southernmost part of its range (Payne 1980; Price et al. 2006). Recent expansion into the northeastern United States has raised concerns about its potential to impact pine species beyond its historical range, including pitch pine (*P. rigida*) and red pine (*P. resinosa*), and which inhabit ecosystems that are already threatened. Forest stands with a higher proportion of SPB host tree species are generally more susceptible to infestation and damage (Aoki et al. 2018).

Economic losses caused by SPB have been substantial, reaching an estimated 1.7 billion USD between 1990 and 2004. This high impact is largely because SPB’s primary hosts (loblolly and shortleaf pine) are among the most economically valuable tree species in the southeastern United States (Pye et al. 2011). Furthermore, SPB outbreaks have strongly impacted ecosystems (Ciesla 2011). Historically, SPB outbreaks recurred every 5 to 7 years in the southeastern United States. However, since 2000, the extent of affected areas in the Southeast has declined significantly. At the same time, isolated populations and outbreaks have emerged in the northeastern United States, in New Jersey, New York (Long Island), Connecticut, New Hampshire, and Maine that had previously not experienced SPB outbreaks (Kanaskie et al. 2023).

The northern range limit of SPB is strongly constrained by cold temperatures in winter. Its population growth rates sharply decrease in areas that experience temperatures below -16°C (Ungerer et al 1999; Lombardero et al. 2000) making SPB one of the least cold tolerant bark beetle species in North America. To assess changes in climate suitability across the SPB range, we applied the model developed by Lesk et al. (2017), which uses extreme winter cold events as a proxy for overwintering survival. This model incorporates a linear Newtonian heat flux framework to estimate phloem temperatures, where SPB eggs are deposited and larvae develop (Trân et al. 2007; Lesk et al. 2017). We selected the survival proportion of individuals (*Psurv*) as the performance indicator for SPB. This variable is calculated based on the temperatures experienced by each life stage, as cold tolerance varies across developmental stages (Wagner et al. 1984; Lombardero et al. 2000; McManis et al. 2018). The model estimates the frequency and intensity of lethal temperature exposures using phloem temperature as a proxy. Although SPB also occurs in parts of the southwestern United States, including Arizona, our analysis was restricted to eastern North America, where the species has historically caused the most significant damage (Dodds et al. 2018; Fettig et al. 2022).

### Spruce beetle (Dendroctonus rufipennis)

Spruce beetle (SB) is a native bark beetle distributed throughout the boreal and subalpine spruce forests of northwestern North America although it is also present in spruce forests of northeastern North America (Holsten et al. 1999). It primarily infests weakened, mature and overmature Engelmann spruce (*Picea engelmannii*) and white spruce (*P. glauca*). Spruce beetle outbreaks can be initiated by factors such as favorable climatic conditions, windthrow, or drought, and may lead to extensive tree mortality across large landscapes. When populations reach high densities, the beetle is capable of successfully attacking and killing healthy, vigorous trees. Its activity alters forest structure, composition, and creates fuels for subsequent fire disturbance events (Bleiker and Brooks 2021).

Under cool climatic conditions, SB typically requires two years to complete its life cycle (Bleiker and Brooks 2021). However, anomalously warm temperatures can accelerate development, enabling individuals to complete their life cycle in a single year. This temperature-driven shift toward a univoltine (one-year) life cycle is called adaptive seasonality and significantly increases population growth rates and outbreak potential. We used a model based on Hansen et al. (2001) to predict the probability of SB to complete its life cycle in one year (*PropUni*). The binomial model is based on accumulated hourly temperature above 17°C 40 to 90 days after peak flight of the adult stage. This model predicts that the proportion of univoltine broods increases with accumulated temperatures, making the model’s predictions sensitive to the cold constraint of the species. Although SB occurs across the range of spruce (*Picea* spp.) in North America, we restricted our analyses to the western portion of its range, where its impacts are greatest.

### Emerald ash borer (*Agrilus planipennis*)

Emerald ash borer (EAB) is a non-native, wood-boring beetle from East Asia that was first detected in North America in the early 2000s near Detroit, Michigan and Windsor, Ontario (Haack et al. 2002), although it was likely introduced in the 1990s (Siegart et al. 2014). Since then, it has spread rapidly across much of eastern North America, causing near-total mortality of almost all native North American species of ash trees (*Fraxinus* spp.) in affected areas. The potential cost of this exotic species in Canada has been evaluated around 1.2 to 1.4 billion CAD (Hope et al. 2021). Larvae feed on the phloem and cambium, disrupting nutrient transport and leading to tree death within a few years. The species poses a major threat to urban forests, riparian systems, and native ash populations. Of particular importance is black ash (*Fraxinus nigra*) which is a keystone cultural species that holds cultural and practical significance for First Nation and Indigenous communities in North America, its decline could have direct impacts on their traditional practices and livelihoods (Siegert et al. 2023).

The EAB is a freeze-intolerant species (Crosthwaite et al. 2011) that demonstrates high phenological plasticity to extreme winter cold (Duell et al. 2022). Southern populations of the insect appear to be constrained by temperatures below -30 °C (Crosthwaite et al. 2011) but this can range to as low as -50 °C in the northern part of the range (Duell et al. 2022), allowing them to survive much more severe cold snaps. We used the model mortality rate estimates (Mortality) as the performance indicator for EAB. This variable is based on winter temperatures beneath the bark of ash trees, using a Newtonian cooling approach to simulate under-bark thermal minima during cold events (Vermunt et al. 2012). Although EAB was introduced recently, we computed model outputs for the entire 1951–2022 period. This approach allowed us to evaluate the historical evolution of climate suitability for EAB prior to its arrival in North America and in the majority of its present range, serving as a retrospective baseline to understand how favorable conditions may have developed, and to anticipate how they may continue to evolve in the coming decades under ongoing climate change. Emerald ash borer also now occurs in western North America (MacQuarrie et al. 2025a) but we restricted our analyses to the eastern portion of its range where it was first introduced, and its impacts have been greatest.

### Climate data

To evaluate each ecophysiological variable performance for all species, we used daily temperature data from the ERA5 reanalysis dataset (Hersbach et al. 2020). Developed by the European Centre for Medium-Range Weather Forecasts (ECMWF), ERA5 offers high-resolution global climate data by assimilating historical observations with state-of-the-art numerical modeling (Hersbach et al. 2020). The dataset provides hourly climate variables, including atmospheric, oceanic, and land surface conditions, dating back to 1940, with a horizontal resolution of approximately 31 km (0.25° x 0.25°). For our analyses, we extracted daily temperature records from 1950 to 2022, limited to grid cells covering the United States and Canada. These data were then processed using BioSIM11 (Régniere 2017) to drive each ecophysiological model for all pest species separately.

### Host tree datasets

The distribution and abundance of host tree species vary spatially across North America, and climate-driven shifts in pest suitability are expected to influence the extent of host biomass at risk. Due to the absence of a comprehensive, standardized raster dataset for tree species biomass at the continental scale, we used national datasets for Canada and the United States separately. For Canada, host tree biomass was derived from the National Forest Inventory of Canada (http://nfis.nfi.org, NFI 2023), which provides species-specific aboveground biomass estimates based on a k-nearest-neighbor (kNN) interpolation of ground-based photo plot data, combined with MODIS-derived remote sensing imagery. This dataset offers 250 m resolution biomass estimates for each tree species occurring in Canada for the year 2011.

In the United States, host species biomass was obtained from the Forest Inventory and Analysis (FIA) BIGMAP Tree Species Aboveground Forest Biomass dataset (Wilson et al. 2018). This dataset provides tree species biomass estimates for the conterminous United States at a 30 m resolution, based on Landsat 8 OLI imagery for the 2014–2018 period. Similar to the Canadian approach, biomass estimates were derived through kNN interpolation using FIA survey plots. As the dataset excludes Alaska, subsequent analyses of host biomass were limited to the conterminous United States.

For both datasets, species-specific biomass values were aggregated to the ERA5 grid-cell scale. For each pest species, the total biomass of all important host species was summed within each ERA5 grid cell to produce a standardized map of host biomass at risk for all of North America. Tree species considered as hosts are listed in Supp. Mat. S1 for each pest species.

## Analyses

### Trends in weather-driven pest performance indicators

To characterize weather-driven pest performance trends we extracted daily temperature data from year *t* in the ERA5 dataset. For species developing over two calendar years, we also used daily weather data from *t* – 1. Weather data were then integrated into BioSIM11 to simulate the annual development of each pest species at the grid-cell level, using the corresponding species-specific ecophysiological model. Performance indices were then compiled annually from these simulations.

To assess temporal trends, we applied the Thiel–Sen slope estimator to the 1951–2022 time series for each ecophysiological performance indicator at the grid-cell level. The Thiel–Sen method, being non-parametric and robust to outliers, is well suited for detecting climate trends (Huth and Pokorná 2004). Prior to trend analysis, the time series were pre-whitened to remove temporal autocorrelation so as to preserve independence of data. Trend significance was evaluated at *P* ≤ 0.1 using the *tfpwmk* function from the *modifiedmk* package v1.6 (Patakamuri and O’Brien 2021) in R v4.3.2 (R Core Team 2022).

We mapped spatially significant trends for all variables to highlight regional patterns. Given that climate change since 1950 could vary along a west-east gradient within pest species range (Bush and Lemmen 2019), we further analyzed longitudinal patterns by averaging Thiel–Sen slope values across grid cells sharing the same longitude. Nonsignificant trends were assigned a slope of zero.

To focus the analysis on areas where each pest species is known to occur, we restricted our assessment to grid cells falling within the current distribution range of each species. Continent- wide boundary’ layers of official ranges are difficult to obtain for most pest species. We obtained range maps for ESBW (Régnière et al. 2012), WSBW (Régnière and Nealis 2019), and MPB from previously published sources. For SB, the western range was delineated based on the distribution of spruce species within major watersheds draining into the Pacific Ocean, reflecting the beetle’s close association with spruce-dominated forests (Bleiker and Brooks 2021). For the remaining species, in the absence of official or standardized range maps, we relied on the iNaturalist Open Range Map Dataset (iNaturalist 2025). This dataset is derived from hundreds of thousands of geo-referenced community-contributed observations and provides interpolated species range maps based on spatially implicit neural network models. Maps are generated through spatial interpolation using deep learning models based on Spatial Implicit Neural Representations (Cole et al. 2023). According to species experts, these maps offer a reliable approximation of current species distributions across North America. Trends were reported per decade as well as over the full 1951–2022 period.

### Assessing latitudinal and elevation shifts

We further assessed spatial shifts in areas characterized by highly suitable climate conditions. That is, those areas where weather-driven performance indices were greatest. In the absence of species-specific thresholds, “high suitability” was operationally defined as when the annual performance values exceeded the median across all years within the species’ current range.

Using Thiel–Sen-derived estimates, we predicted species’ performance for both 1951 and 2022 and then identified areas classified as highly suitable in each year. We quantified spatial shift by estimating the southern, mid- and northern predicted latitude where conditions were highly suitable in both years, by longitude. These areas were respectively calculated as the 5th, 50th and 95th percentile of latitude showing predicted suitable values for each longitude in 1951 and then in 2022 (see Boulanger et al. 2025 for a similar approach). This approach allowed us to assess whether highly suitable climatic conditions at the northern and southern range margins are shifting in a manner consistent with those in the core of the species’ suitable range over the 72-year period. For presentation, the positions (5^th^, 50^th^ and 95^th^ percentiles) of suitable conditions were reported by longitudes and grouped into four geographic regions: Western (>120°W), Central (120°–95°W), Eastern (95°–64.5°W), and Atlantic (<64.5°W).

For species primarily occurring in mountainous regions, namely SB, MPB, and WSBW, we also analyzed shifts in elevation. Elevational shifts at lower, mid-, and higher elevations (5^th^, 50^th^, and 95^th^ percentiles, respectively) were assessed using the same methodology as for latitudinal shifts. Here, however, we analyzed shifts along a latitudinal (south to north) gradient instead of longitudinal, to determine whether elevation shifts varied with latitude. This approach was taken because these species are primarily restricted to western North America, where their distribution spans a limited longitudinal range.

We emphasize that the models used to estimate performance indicators for WSBW, SM, and ESBW capture constraints at both the cold and warm edges of each species’ climatic niche. In contrast, the models applied to SB, HWA, MPB, SPB and EAB primarily represent limitations at these species’ colder range margins, with limited sensitivity to warm-edge constraints. As a result, northward or elevational shifts in highly suitable conditions must be interpreted differently across these groups. For SB, WSBW, and ESBW, changes in the 5th, 50th, and 95th percentiles of suitability reflect shifts across the full range of climatic suitability, i.e., from cold-limited northern margins to warm-limited southern margins. Conversely, for HWA, MPB, SPB and EAB, changes in the 5th, 50th, and 95th percentiles of suitability reflect changes in the cold-limited portion of the distribution, i.e., the expansion of the northern (or high-elevation), mid-, and southern (or low-elevation) boundaries of cold constraints.

### Trends in host biomass exposed

Using the ERA5 grid cells identified as having highly suitable climatic conditions for each pest species, we computed the total annual host tree biomass within these areas. For each year, and for each pest species individually, we summed the biomass of host trees within grid cells classified as highly suitable, yielding an annual estimate of total host biomass "exposed" to pests. To assess temporal trends in exposed host biomass over the 1951–2022 period, we applied the same approach described above for the performance indicator, using the modified Thiel–Sen slope estimator. Trends were estimated for each pest species separately.

All analyses were performed on R v4.3.2 (R CoreTeam 2022).

## Results Overall trends

Four out of eight species exhibited significant temporal shifts in climate-driven performance indicators as averaged over their current ranges (Fig. 1). Overall performance indicators significantly (P < 0.1) increased for SM, HWA, and SB, decreased for EAB, and remained stable for all other species (ESBW, WSBW, MPB, SPB). Yet, spatial patterns of change varied across species’ ranges (Fig. 2). In general, performance tended to increase in the northernmost portions of species’ ranges and/or at higher elevations. An exception was SM, for which performance remained relatively stable across its entire range over the study period, except at its northernmost margin. Unlike the other species, HWA showed the greatest increase in performance in the southern part of its range, while increased performance northward was more limited. Performance in the southernmost parts of most other species’ ranges remained stable or declined. For instance, climate conditions became less favorable for ESBW, WSBW, and SPB in their southern extents. Substantial changes in performance were also detected outside of current species’ ranges, with increasing performance frequently occurring north of the current range of the species; however, these areas were excluded from the core analyses.

**Fig. 1.**
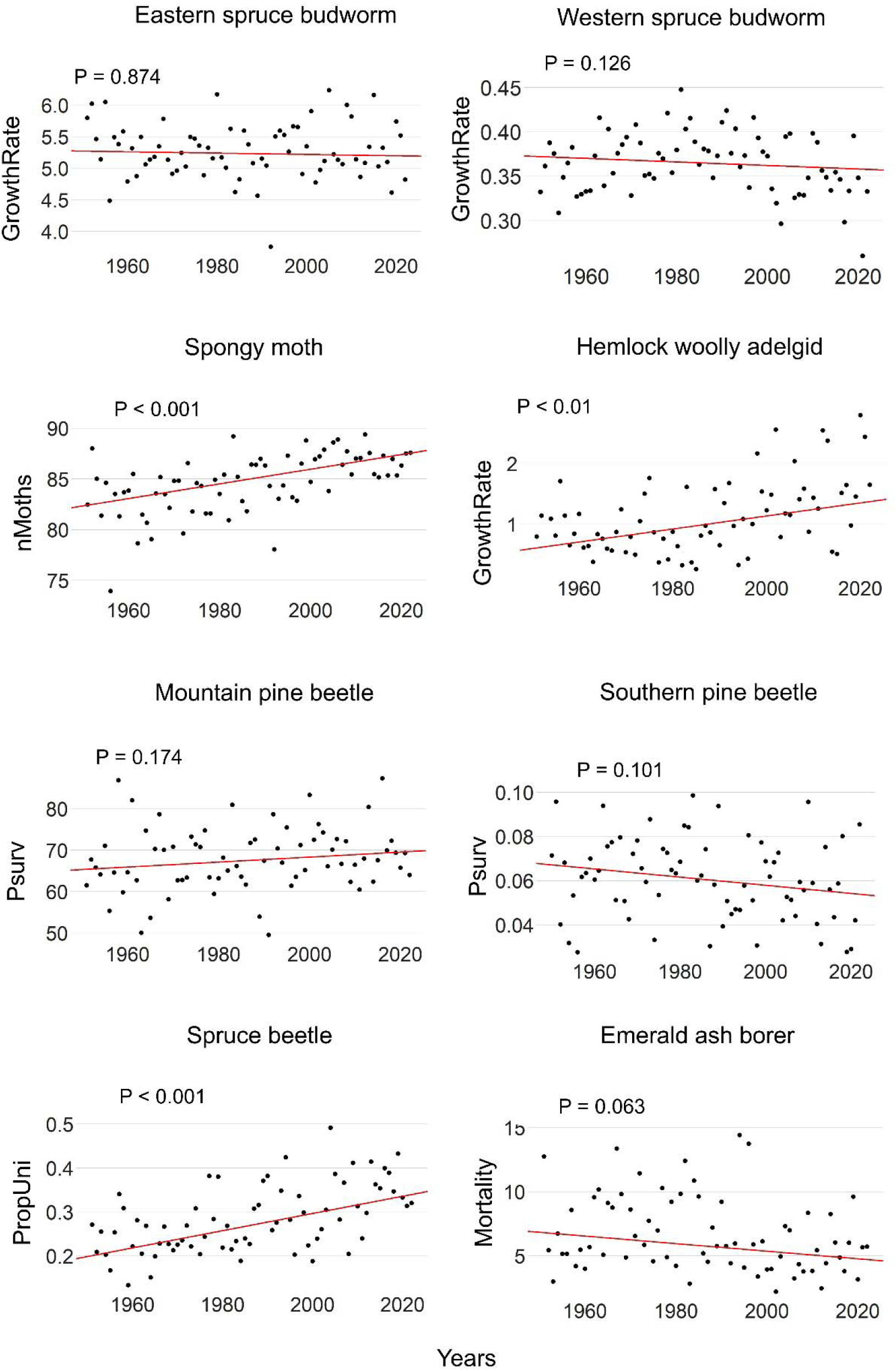
Trends in weather-driven performance indicators as averaged over the species range for the eight North American forest insect species considered in the present study. The red line represents the Sen slope linear trend.

**Fig. 2.**
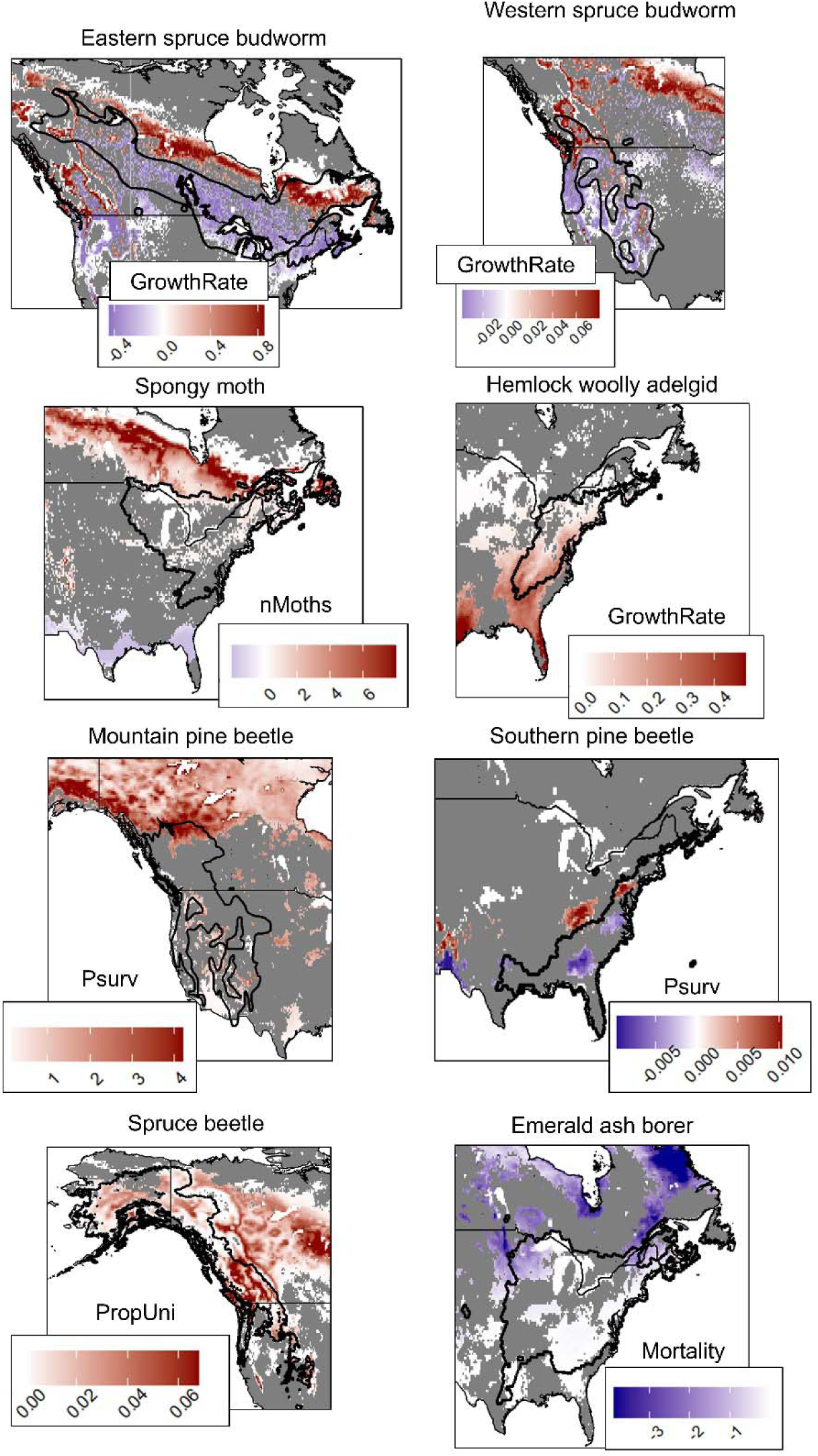
Maps showing decadal trends in weather-drive performance indicators for eastern spruce budworm and western spruce budworm, spongy moth, hemlock woolly adelgid, mountain pine beetle, southern pine beetle and spruce beetle, and emerald ash borer. Trends are expressed at a decadal time scale and calculated for the 1951- 2022 period at the ERA5 cell resolution (∼31 km) across North America. ERA5 cells showing non-significant trends (P-value > 0.1) appear in gray. Species range is shown by the black contour line. GrowthRate: predicted population growth rates; Psurv: probability of survival; nMoths: maximum number of adults; PropUni: proportion of univoltinism; Mortality: mortality rates.

Overall, temporal trends in species performance revealed a general northward shift in areas classified as highly suitable for all pest species (Fig. 3). In most cases, northward displacement was more pronounced at the northern and mid-range portions of the highly suitable area, while shifts at the southern margins were comparatively limited. The most substantial northward shifts (∼60 km on average) at the northern and mid-range boundaries were observed for ESBW and EAB. Moderate shifts (∼40 km) were simulated for SM. Western spruce budworm was the only species to show substantially greater northward shifts for mid-range (97.1 km) and southern (84.8 km) boundaries than for the northern boundary (10.8 km). Other species exhibited more limited climate range adjustments, generally averaging below 40 km (Fig. 3).

**Fig. 3.**
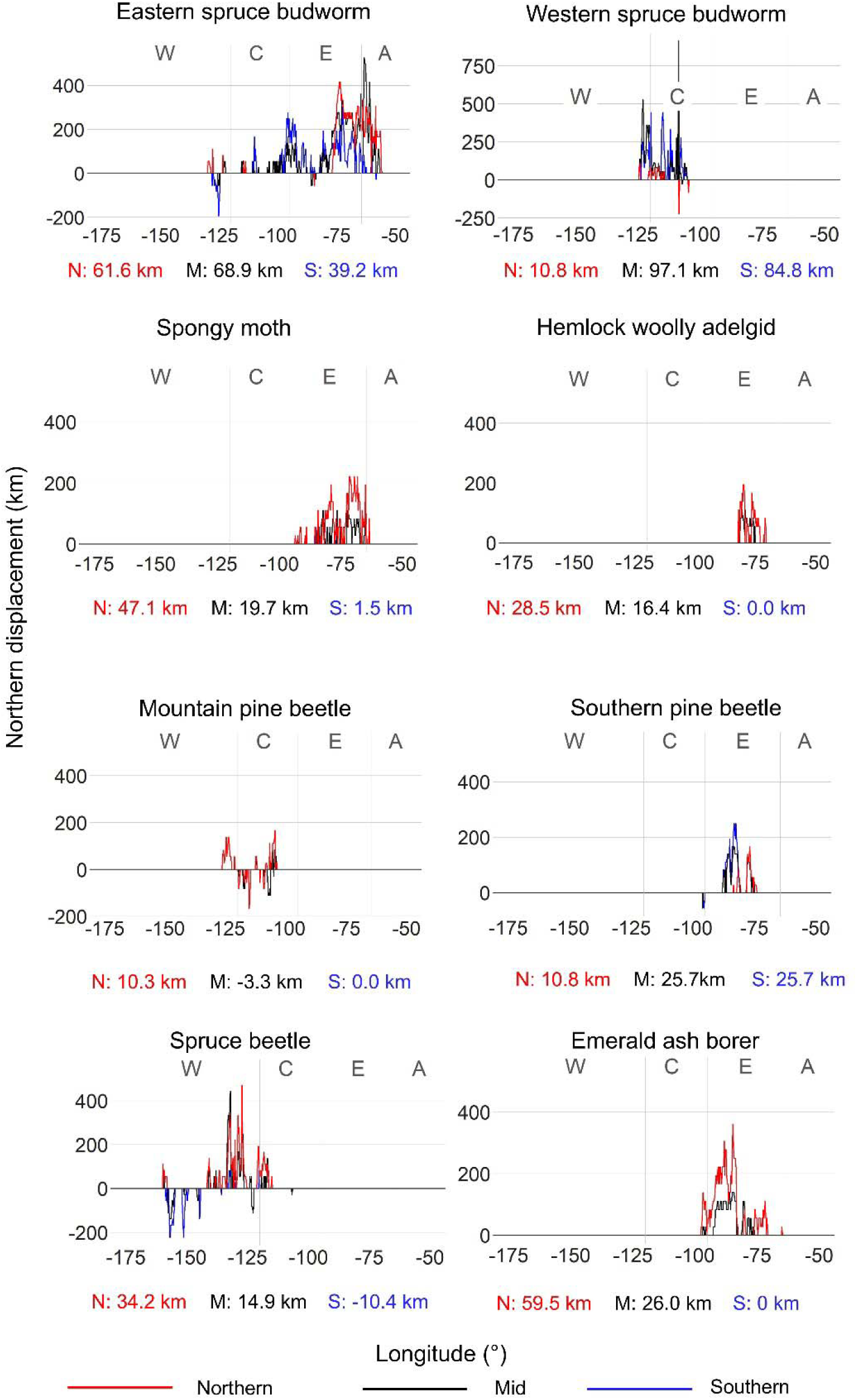
Northern displacement (in km) for the northern (red), mid-range (black) and southern (blue) limit of highly suitable areas for each species as averaged by longitude within the pest species range between 1951 and 2022. West, Central, East and Atlantic regions are arbitrarily delineated using the following longitudes: -120°, -95° and - 64.5° respectively. Mean northward displacements for the northern, mid-range and southern boundaries across pest species range are also shown under each species graph.

Locally, shifts in highly suitable conditions, especially along the northern margins, are much more pronounced than when looking at averages (Fig. 3). For example, the northern edge of highly suitable conditions for ESBW shifted between 200 and 400 km in the Eastern and Atlantic zones, whereas shifts were more modest (< 200 km) in the Western and Central zones. Similar northward displacements, approaching or exceeding 200 km, were also observed for the northern margins of the highly suitable conditions for SM, HWA, SPB, and EAB, primarily within the Eastern zone. For species primarily distributed in western North America, latitudinal displacement patterns were more complex, likely due to topographic variability associated with mountainous regions. In the case of WSBW, northward shifts were greatest in the Western zone, ranging from 200 to > 500 km, while more limited displacements (< 200 km) were observed in the Central zone with very few exceptions (Fig. 3). MPB showed a variable pattern, with shifts of < 200 km northward in the Western zone, followed by alternating southward and northward displacements across its range moving eastward. Finally, for SB, the westernmost portion of its highly suitable area sometimes shifted > 150 km southward, whereas areas located east of approx 140°W exhibited northward shifts ranging from 200 to > 400 km (Fig. 3).

## Elevational trends

Among the three species primarily distributed in the mountainous regions of western North America, both WSBW and SB exhibited significant upward shifts in elevation of highly suitable climate conditions. No consistent elevational shift in cold constraints was observed for MPB (Fig. 4). For SB, the most pronounced changes occurred at the upper elevation boundaries, with average increases of +88 m. In contrast, shifts at mid and lower elevation boundaries for SB were comparatively limited. Elevational shifts also varied along a latitudinal gradient. The greatest upward displacements were observed at the highest elevation margins in the northern half of the species’ ranges, where shifts > +200 m at several latitudes (Fig. 4). For WSBW, elevational shifts were slightly different than for SB, with greatest elevational shifts occurring at mid- and lower elevation, and mostly between 35 and 45°N. Largest elevational shifts at higher altitude occurred at mid-latitude (37 and 41°N) but also at more northerly latitudes (> 47°N)

**Fig. 4.**
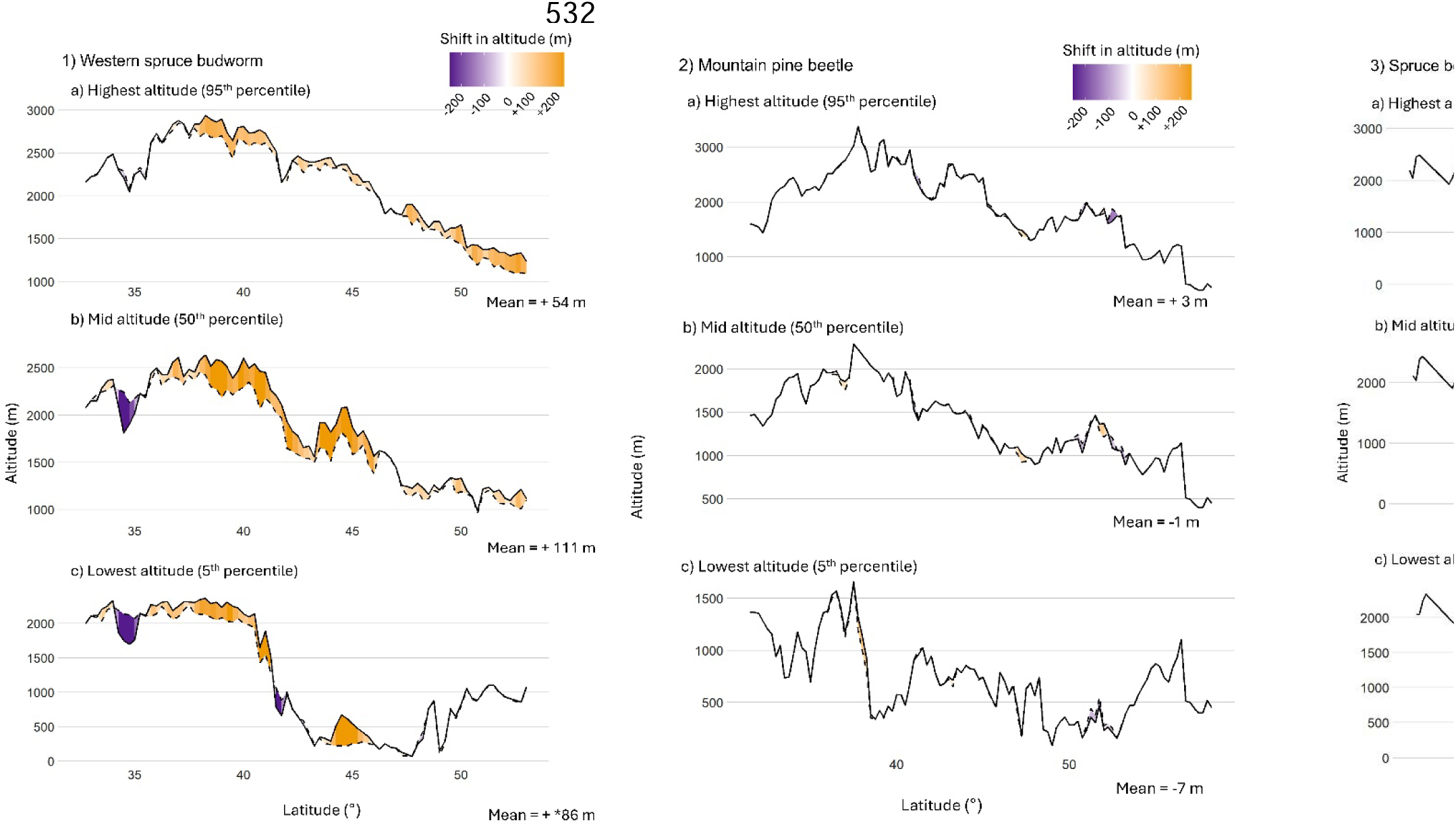
Altitudinal location predicted according to the trend analyses in 1951 (dotted line) and in 2022 (solid line) for the a) highest altitude (95th percentile), b) mid altitude (50th percentile) and c) lowest altitude (5th percentile) limit of highly suitable areas for the three mountainous species,e. 1) western spruce budworm, 2) mountain pine beetle and 3) spruce beetle, as averaged by latitude within the pest species range. Mean altitudinal displacements for the highest, mid and lowest altitude boundaries across pest species range are also shown under each species graph.

## Trends in area having highly suitable conditions and in host biomass

The extent of areas classified as having highly suitable climate conditions significantly increased for five of the eight pest species examined, while no significant trends were observed for the remaining three (Fig. 5). The most pronounced increases were observed for HWA and SB, with gains of +63% and +55%, respectively. More moderate, yet still substantial, increases in suitable area were recorded for SM, and EAB, ranging from +40% to +32%. WSBW was the only species showing a decrease in highly suitable areas within its range. Consistent with changes in area, the total host biomass exposed to highly suitable conditions also increased for most species between 1951 and 2022 (Fig. 6). Exposure was particularly dramatic for HWA and SB, with a nearly +230% and +73% increase, respectively in total biomass at risk. Other species, including EAB (+42**%)**, and SM (+33%), experienced important increases in biomass exposure, while exposure remained relatively stable for the remaining species.

**Fig. 5.**
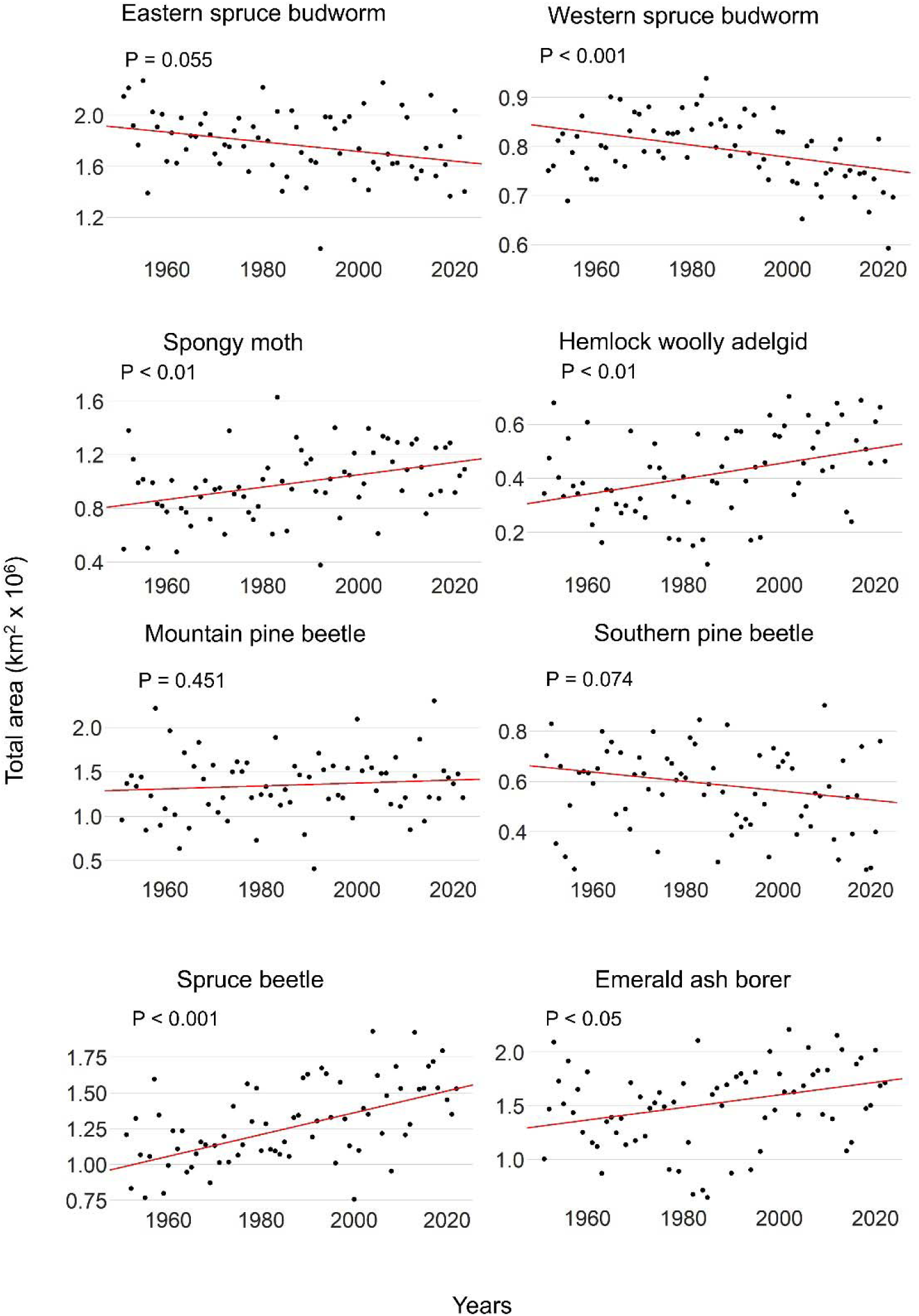
Trends in the extent of areas classified as highly suitable based on yearly climate conditions for each pest species from 1951 to 2022. The slope of the trend (red line) was estimated using the Thiel–Sen method, and associated *P*-values are shown to indicate statistical significance.

**Fig. 6.**
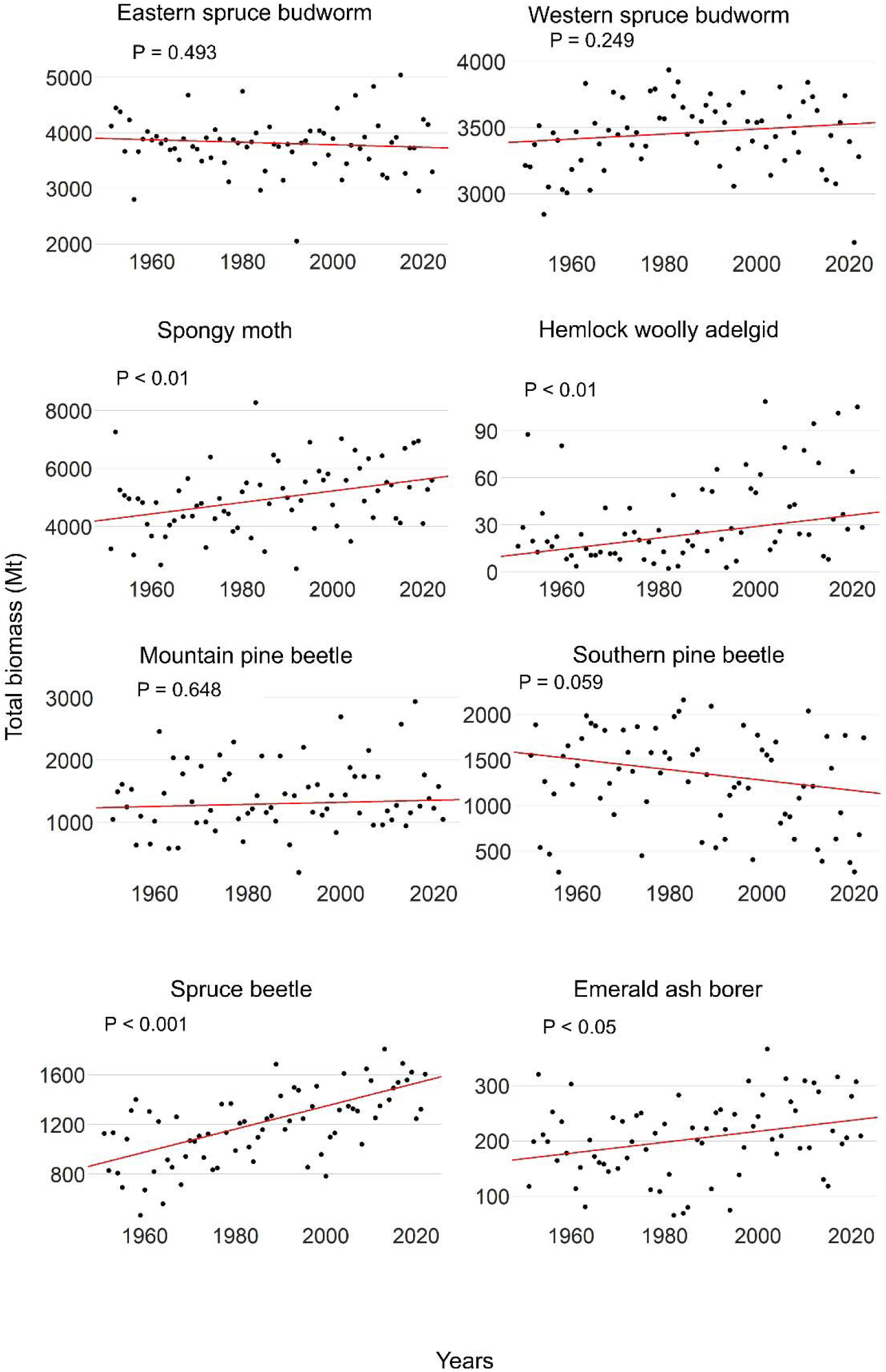
Trends in total host biomass (in megatons) located within areas classified as highly suitable based on yearly climate conditions for each pest species from 1951 to 2022. Thiel–Sen slopes (red lines) are presented along with corresponding *P*-values to indicate the significance of observed trends.

## Discussion

Our analyses reveal substantial changes in climate suitability since 1950 resulting in pronounced northward and elevational shifts for most pest species. We also show that the magnitude and spatial patterns of these shifts vary both geographically and among species. Heterogeneous shifts in the climatically suitable range of pest species are primarily driven by how each species responds to limiting climate constraints to climate change. This highlights the importance of selecting appropriate suitability metrics tailored to each species’ ecological and physiological sensitivities. Overall, increasingly favorable conditions for some species at northern latitudes and higher elevations reflect a relaxation of climatic constraints at the cold edge of species’ distributions. These constraints are primarily driven by the improved capacity to complete the life cycle (e.g., SM, ESBW, WSBW, SB) and increased survival during the cold season (e.g., MPB, SPB, HWA, EAB). In contrast, we also show that recent warming has reduced the climatic suitability at the southern edges for forest insect species for which warmer temperatures can be detrimental, notably during diapause (e.g., ESBW & WSBW, see: Régnière et al. 2012; Régnière and Nealis 2019; Boulanger et al. 2025) or during the developmental season (SM; see Tobin et al. 2014). Furthermore, topographic and geophysical constraints, along with spatial heterogeneity in climate trends, can lead to regionally divergent responses in species range shifts, even within the same species. For instance, the more pronounced northward shift of ESBW in eastern Canada may be attributed to improving climatic suitability within its current distribution in this region. In contrast, in western Canada, climatic improvements have largely occurred beyond the species’ historical range limits, resulting in more constrained measurable shifts (Boulanger et al. 2025). Likewise, asymmetry in recent upslope shifts is partly attributable to geographic and topographic constraints that differ between southern and northern regions. In southern areas, e.g., the southwestern United States, upward range shifts are limited because species are already near the highest available elevations (∼2,500-3,000 m asl), leaving little to no room for further upslope expansion. In contrast, northern regions often present greater elevational room near or above the historical colder boundary (e.g., in the Canadian Rockies), allowing cold-range boundaries for species like WSBW and SB to continue migrating upslope in response to warming.

The range shifts observed in this study are consistent with previous findings. For instance, the range of MPB expanded northwards by nearly 400 km between 1965 and 1996, driven by warming temperatures (Trân et al. 2007; Sambaraju et al. 2019). While our study did not detect a significant elevational shift in cold constraints (see the Limitations section), evidence suggests that MPB has adapted to higher elevations through shifts in the timing of events in its life cycle enabled by pronounced winter warming that has relaxed thermal limits on beetle survival and development (Hicke et al. 2006; Weed et al. 2015). Likewise, previous studies have shown that in recent decades the isoline for lethal winter temperatures causing 50% mortality in SPB populations has shifted approximately 1.5° latitude northward (about 165 km) (Ayres and Lombardero 2000). Simulated shifts in climatic suitability in the present study are also consistent with observed changes in pest population dynamics and outbreak distributions. For example, Pureswaran et al. (2018) reported that the most recent ESBW outbreak in eastern Canada occurred significantly farther north than previous outbreaks. Our results show that this region corresponds to the one that experienced the most important increase in climate suitability for ESBW within its range. Concurrently, regions in the southern portion of the ESBW range are yet to be affected by the current outbreak. The location of these regions aligns with areas in our analyses where simulated growth rates have significantly declined over the past seven decades (Boulanger et al. 2025). Similarly, long-term declines in WSBW outbreak frequency in the warmer regions of its distribution have been linked to decreasing climatic suitability (Alfaro et al. 2018; Tai and Carroll 2022), a pattern that is corroborated by our simulations. Our findings align with recent observations indicating that the 1996–2016 WSBW outbreak in British Columbia shifted northward and upslope (Maclauchlan et al. 2018, Régnière and Nealis 2019, Srivastava et al. 2024). Furthermore, warming temperatures at higher elevations have been associated with increased frequency and severity of MPB and SB outbreaks in the western United States and western Canada ( Berg et al. 2006 ; Bentz et al., 2016). Milder winters have enhanced MPB and SB survival and reproduction at higher elevations and northern latitudes, contributing to recent unprecedented outbreaks in previously unsuitable regions (Goodsman et al. 2019; Berg et al. 2006; Chavardes et al. 2012). Our results thus suggest that recent climate warming has facilitated both phenological adaptation and outbreak potential in previously climatically constrained areas. This also aligns with a growing body of global evidence indicating that climate change over the past several decades is driving forest insect pests and the severity of their impacts to expand poleward and upslope across multiple ecosystems and continents (Jepsen et al. 2008; Yasyukevitch et al. 2015).

We showed that shifts in highly suitable conditions are more important at the colder edge (either northern or upper elevation) than at the warmer boundaries for several species. This led to range expansion of highly suitable conditions for many species that did not necessarily translate into a similar contraction of suitable conditions at the southern range edge or at lower elevations. The consequence is an increasing area under high climatic suitability. Although the real southern trailing edge could not be effectively modeled for certain species, our simulations nonetheless revealed a more pronounced northward shift in cold-limited constraints at their northernmost margins. More pronounced poleward shifts at high latitudes are likely driven by Arctic amplification, a process in which northern regions warm at rates significantly faster than the global average (Previdi et al. 2021). Similarly, upslope regions are experiencing accelerated warming due to elevation-dependent warming, a phenomenon influenced by mechanisms such as the snow-albedo feedback, spatial heterogeneity in tropospheric warming, and longwave radiation feedback associated with water vapor (Minder et al. 2017). For species where the models do incorporate physiological constraints at the warmer edge of their range (e.g., ESBW, WSBW), range retraction was mostly concomitant with northward displacement. Suggesting that, for these species, the climate envelope shifts might be unaffected by the abovementioned processes.

The climate-driven displacement or expansion of highly suitable conditions, particularly into areas that were previously marginal or unsuitable, indicates an elevated potential for increased risk to forest ecosystems (Cullingham et al. 2011; Burke et al. 2017). Our results clearly indicate that both the total area and the host tree biomass exposed to highly suitable climate conditions have increased for many pest species since the last decades, primarily due to the expansion and northward or elevational shifts in the climatic suitability ranges of these species. For instance, the recent increase in climate suitability for ESBW in the north has heightened black spruce (*Picea mariana*) vulnerability by improving pest–host synchrony and increased the risks of mortality and ecosystem shifts (Pureswaran et al. 2015, 2018). Warming temperatures at the expanding northern edge of the MPB range have been shown to alter the beetle’s phenology which increases the likelihood of synchronized mass dispersal events and heightening the risk of outbreaks (Bleiker and van Hezewijk 2016) and further exposing naive populations of lodgepole pine to MPB attack (Cudmore et al. 2010; Cullingham et al. 2011; Renwick et al. 2016; Burke et al. 2017). Recent northward and elevational expansions in climate suitability we simulated are also putting ecologically sensitive ecosystems at increased risk. For instance, rising climate suitability near the northern edge of SPB range is now threatening pitch pine (*Pinus rigida*) populations, a species of limited distribution and ecological importance in that region. Likewise, climate-driven northward expansion of HWA in recent decades has increasingly threatened eastern hemlock (*Tsuga canadensi*s) stands, a keystone species of climax forest communities in northeastern North America (Paradis et al., 2008), triggering significant changes in forest composition and structure. Furthermore, increased suitability at higher altitude increases the vulnerability of high-elevation five-needle pines, including whitebark pine (*Pinus albicaulis*), limber pine (*Pinus flexilis*), and bristlecone pine (*Pinus longaeva*), to MPB outbreaks (Buotte et al. 2016; Howe et al. 2021; Tomback et al. 2021; Macfarlane et al. 2023; Bentz et al. 2022; Airey and Taylor 2024). Rising climate suitability at the northern edge of EAB range is threatening black ash (*Fraxinus nigra*), a culturally vital species for many First Nations, with potential consequences for traditional practices and cultural continuity (Siegert et al. 2023). In this context, implementing targeted adaptation strategies is becoming increasingly critical to enhance the resilience of forest stands against the climate-driven rise in forest pest suitability across North America.

Shifts in climate suitability for forest insect pests pose growing challenges to existing management frameworks and carry important implications for the forest industry, particularly in regions where key host species have high economic value. The intensifying pressure on black spruce in the northern range of ESBW has potentially significant economic implications as this tree species is a key roundwood resource in Canada (Natural Resources Canada 2025). In addition, the last MPB outbreak in western Canada, driven by rising climatic suitability as we showed, caused extensive pine mortality which resulted in large-scale salvage logging, leading to long-term declines in wood quality and market stability (Dahr et al. 2016). Increased salvage wood from pest outbreaks could shift towards the wood pellet industry, presenting both sustainability challenges and opportunities to bolster bioenergy supply, if managed within ecological and climate objectives (Neidermeier et al., 2020). The forest sector may need to adapt to this emerging reality by developing more flexible supply chains, improved forecasting tools, and policy frameworks that account for both increasing climate-driven forest pest impacts and volatile timber markets (Halofsky and Peterson 2016).

By analyzing multiple pest species simultaneously, we identified widespread northward and elevational shifts in climate suitability that could increasingly expose previously unaffected forest ecosystems to overlapping, potentially cumulative and interacting, biological disturbances. This is particularly evident in western Canada, where the regions with higher climatic suitability for WSBW, MPB, and SB are expanding. Such multispecies shifts could increasingly allow for the geographic overlap of several pest species in areas where none or few of these disturbances have historically occurred. This overlap could lead to novel combinations of pest pressures, with potentially significant ecological consequences. Furthermore, increasing climatic suitability for forest pests may intensify their interactions with other disturbance agents, potentially compounding their ecological and economic impacts. By altering forest composition and structure, pest-driven mortality can modify fuel loads and continuity, thereby affecting wildfire behavior and severity (Loehman et al. 2017; Keane et al. 2022; Romualdi et al. 2023; Woo et al. 2024). Additionally, novel synergies between insect outbreaks and other climate-driven disturbances, such as drought or windthrow, may exacerbate widespread host mortality, particularly in vulnerable ecosystems (Wong and Daniels 2016). These cascading disturbances in novel habitats may increase the susceptibility of future forests to additional abiotic and biotic stressors under continued anthropogenic climate forcing (Perovich and Sibold 2016).

Our results indicate that climate change has already contributed to the increasing climatic suitability and geographic spread of exotic forest pest species in North America since the mid- 20th century. Specifically, all three non-native species examined (SM, EAB,HWA), have experienced a significant expansion in the extent of highly suitable climatic conditions, particularly at the northern margins of their current ranges (Sharov et al. 1999; Morin et al. 2009; Fitzpatrick et al. 2012). These findings are consistent with field observations showing that all three species have continued to expand their ranges over recent decades, particularly into more northerly regions (e.g., Cornelsen et al. 2024; MacQuarrie et al. 2025b). This has resulted in a marked increase in both the area at risk and the total host biomass exposed to highly suitable conditions for these exotic pests. An increase in host volume at risk can create positive feedback loops by supporting larger exotic pest populations, thereby accelerating their spread and intensifying outbreak dynamics (Morin et al. 2009). Our simulations reveal that the observed northward expansions of exotic pest species, mirroring patterns seen in native species, are not uniform but instead reflect distinct, species-specific responses to climatic variables. These divergent responses underscore the varying physiological sensitivities of exotic species to temperature and other climatic drivers.

Our analysis suggests that the recent northward expansion of several exotic forest insect species is not solely attributable to dispersal dynamics but is also likely facilitated by increasingly favorable climatic conditions. Although the spread of these exotic species is shaped by complex interactions among human-mediated dispersal and ecological connectivity (Sharov et al. 2002; Baranchikov et al. 2024; MacQuarrie et al. 2025b), results suggest that recent climate change may have facilitated their continued northward expansion (Régnière et al. 2009). For instance, although EAB was introduced only in the 1990s–2000s, it has rapidly spread across eastern North America, especially in areas dominated by *Fraxinus* species (Liang and Fei 2014). Our simulations indicate that the Great Lakes region, its presumed point of introduction, has experienced one of the largest climate-driven declines in winter mortality (Figure 2), likely facilitating EAB’s establishment and spread. In other words, had EAB arrived several decades earlier, its northward expansion would likely have been more limited due to colder and less suitable climatic conditions (MacQuarrie et al. 2019). Similarly, the recent northern expansion of SM (Régnière et al. 2009) has occurred in areas where climate suitability increased most markedly within its current distribution. Yet, a notable and somewhat unexpected result was the marked increase in suitability of HWA in its southern range along with a significant improvement in climate suitability at the northern edge. The historically faster spread of HWA in the southern portion of its range could partly be explained by milder winter conditions that rarely reach the critical cold-induced mortality threshold of –1LJ°C, which is known to reduce HWA survival (McAvoy et al. 2017). In contrast, northern regions still regularly experience temperatures below this threshold, constraining HWA’s expansion. Enhanced winter survival in the southern Appalachians, combined with phenological synchrony with eastern hemlock, has likely contributed to a broader and more continuous climatic niche in the south (Evans and Grégoire 2006). Such potential climate-driven expansion of exotic pest species amplifies the potential risks to forest ecosystems by facilitating greater establishment, survival, and spread. Our results thus reinforce the growing body of evidence that climate plays a critical role in facilitating the spread of exotic species, making more area and host volume at risk, underscoring its undeniable influence on invasion dynamics and their inherent impacts on ecosystems, making climate change and invasive forest pests interconnected challenges (Smith et al. 2012).

### Limitations

While the models employed provide reliable estimates of pest performance under cold-limited conditions, their ability to capture warm-edge constraints is more limited. This limitation is particularly relevant for MPB, SPB, EAB, SB, and HWA, where physiological responses to high temperatures (e.g., adaptive seasonality, energy depletion during diapause) are not fully represented in model outputs (Bentz et al. 2010; Mech et al. 2017). Furthermore we stress that the ERA5 reanalysis dataset, although high-resolution on a global scale (∼31 km), lacks the spatial granularity required to accurately capture fine-scale elevational variability, particularly in mountainous regions of western North America.This may partly explain the absence of a detected elevational shift for MPB in our results, although an upward shift reported in previous studies used finer-resolution climate or elevation data (Bentz et al. 2016). Despite this limitation, the models still capture robust and consistent broad-scale elevational and latitudinal trends for WSBW and SB. All species models are primarily temperature-driven, consistent with numerous studies demonstrating that temperature is the dominant climatic factor influencing insect development, phenology, and survival. Yet, it is important to acknowledge that other variables, such as precipitation, can also influence pest performance. In the case of MPB for instance, moisture-related parameters are known to affect host susceptibility and beetle development (Safranyik et al. 2010; Bentz et al. 2010). As such, potential trends in these non-thermal variables may have been overlooked in our analysis. Moreover, our models do not account for potential local adaptation to climate conditions. For example, both EAB and HWA have been reported to exhibit increased cold tolerance at the northernmost extent of their current ranges (Lombardo and Elkinton 2017; Duell et al. 2022). As a result, our simulations may underestimate the potential for range expansion in colder regions where such local adaptation could facilitate a northward expansion. It is also important to stress that these models are strictly climatological and do not account for ecological interactions such as host availability and condition, natural enemies (e.g., parasitoids, predators), or intra- and interspecific competition (Allstadt et al. 2013; Liu et al. 2025). As such, while they offer valuable insight into climate suitability, they do not capture the full complexity of population dynamics or outbreak behavior, which are also shaped by ecological and landscape-level processes. Also, estimates of host volume exposed to suitable climate conditions do not account for temporal changes in host biomass or shifts in host species distribution over time. Thus, these values should be interpreted as indicative of broad trends in potential exposure and not as precise measures of forest vulnerability through time. Remote sensing products are known to underestimate the presence and volume of uncommon tree species at landscape scales. Consequently, the volume of host trees at risk, such as *Fraxinus* spp. for EAB and *Tsuga* spp. for HWA, may be underestimated in our spatial assessments.

## Conclusions

The trends we report in pest species performance indicators over the past seven decades across North America likely represent early signals of continued and potentially accelerating shifts under ongoing anthropogenic climate forcing. Our simulations show that climatic suitability is already increasing beyond the current northern limits of several forest insect pests, including both native and exotic species, suggesting further northward expansion is likely in the coming decades, assuming host availability (Régnière et al. 2009; Smith et al. 2012). These findings are consistent with numerous projections that highlight persistent or intensifying northward and upslope expansions under future warming (Williams and Liebhold 2002; Régnière et al. 2009; Bentz et al. 2010; Régnière et al. 2012; Boulanger et al. 2016). Species constrained at their northern range limits by cold extremes are expected to exhibit particularly strong poleward and upslope expansions due to the rapid warming at high latitudes and elevations (Dukes et al. 2009; Bentz et al. 2016). Although large increases in suitable area may not yet be observed for some species, early signs of northward suitability shifts are likely reflecting the early stage of future climate-driven range expansion (Williams and Liebhold 2002). Yet, realized range shifts will depend on ecological constraints such as the spatial response of host tree species (which migrate more slowly than pests), and the dynamics of natural enemies (e.g., predators and parasitoids; Rodenberg et al. 2024; Régnière et al. 2020, 2021a, 2021b).

The ecological implications of current and future shifts in forest pest species in North America are significant. Given the rapid pace of ecological change, continued monitoring of both native and exotic forest insect pests is essential to detect and document emerging shifts. Early-warning systems are crucial for developing adaptive, proactive forest management and conservation strategies. These efforts will be key to mitigating the cultural, ecological, and economic impacts of expanding pest threats under future climate scenarios (Murdock et al. 2013; Halofsky and Peterson 2016).

## Acknowledgements

The authors would like to thank Adèle DeSaint, Salomon Massoda Tonye, and Maryse Marchand who provided valuable input during the early steps of the research project. Also, authors acknowledge the use of ChatGPT to improve the wording of the manuscript.

## Author’s contributions

**Yan Boulanger:** Conceptualization (lead); Formal Analysis (lead); Methodology (lead); Project Administration; Writing – Original Draft Preparation (lead); Writing – Review & Editing (lead). **Chris MacQuarrie**: Writing – Review & Editing (supporting). **Véronique Martel**: Writing – Review & Editing (supporting). **Jacques Régnière**: Writing – Review & Editing (supporting); Resources (equal); Data Curation (supporting). **Rémi Saint-Amant**: Software (lead); Data Curation (lead); Writing – Review & Editing (supporting). **Kishan Sambaraju**: Writing – Review & Editing (supporting); Resources (equal).

---

**Supplementary material S1.** - Tree species considered as important hosts for each pest species included in this study

| Pests | Host species |
| --- | --- |
| Eastern spruce budworm (ESBW) | *Abies balsamea*<br>*Picea glauca*<br>*Picea mariana*<br>*Picea rubens* |
| Western spruce budworm (WSBW) | *Abies amabilis*<br>*Abies concolor*<br>*Abies lasiocarpa*<br>*Abies grandis*<br>*Picea engelmannii*<br>*Pseudotsuga menziesii* |
| Spongy moth (SM) | All tree species |
| Mountain pine beetle (MPB) | *Pinus albicaulis*<br>*Pinus banksiana*<br>*Pinus contorta*<br>*Pinus ponderosa*<br>*Pinus flexilis*<br>*Pinus monticola* |
| Southern pine beetle (SPB) | *Pinus taeda*<br>*Pinus echinata*<br>*Pinus elliottii*<br>*Pinus virginiana*<br>*Pinus rigida*<br>*Pinus palustris*<br>*Pinus pungens*<br>*Pinus strobus* |
| Spruce beetle (SB) | *Picea engelmannii*<br>*Picea glauca*<br>*Picea mariana*<br>*Picea sitchensis* |
| Emerald ash borer | *Fraxinus americana*<br>*Fraxinus nigra*<br>*Fraxinus pennsylvanica*<br>*Fraxinus quadrangulata* |
| Hemlock woolly adelgid (HWA) | *Tsuga canadensis*<br>*Tsuga caroliniensis* |

